# Detection of IL23p40 via Positron Emission Tomography Visualized Inflammatory Bowel Disease

**DOI:** 10.1101/2022.11.30.518419

**Authors:** Farzaneh Rezazadeh, Nicholas Ramos, Allen-Dexter Saliganan, Najeeb Al-Hallak, Kang Chen, Bashar Mohamad, Wendy N. Wiesend, Nerissa T. Viola

**Affiliations:** Department of Oncology, Karmanos Cancer Institute, Wayne State University, Detroit, MI 48201; Departments of Obstetrics and Gynecology, Wayne State University, Detroit, MI; Department of Gastroenterology, Wayne State University, Detroit, MI; Department of Anatomic Pathology, Corewell Health William Beaumont University Hospital

## Abstract

**Background and aims:** Inflammatory bowel disease (IBD), which includes both Crohn’s Disease (CD) and ulcerative colitis (UC), is a relapsing inflammatory disease of the gastrointestinal (GI) tract. Long term chronic inflammatory conditions elevate patients’ risk for colorectal cancer (CRC). Currently, diagnosis requires endoscopy with biopsy. This procedure is invasive and requires bowel preparatory regimen, adding to patient burden. Interleukin 23 (IL23) plays a key role in inflammation especially in the pathogenesis of IBD and is an established therapeutic target. We propose that imaging of IL23 via immunopositron emission tomography (immunoPET) will potentially lead to a new non-invasive diagnostic approach.

**Methods:** The aim of the present study is to investigate the potential of immunoPET to image inflammation in a chemically induced mouse model of colitis using dextran sodium sulfate (DSS) by targeting IL23 via its p40 subunit with a ^89^Zr-radiolabeled α-IL23p40 antibody.

**Results:** High uptake of the IL23p40 immunoPET agent in mice were displayed in DSS-administered mice, which correlated with increased IL23p40 present in sera. Competitive binding studies confirmed the specificity of the radiotracer for IL23p40 in the GI tract.

**Conclusion:** Taken together, these promising results set the stage for developing this radiotracer as an imaging biomarker for IBD diagnosis. Noninvasive imaging of IBD with IL23p40 immunoPET may help physicians in their treatment decisions for IBD management.

## Introduction

Inflammatory bowel disease (IBD) is represented by a chronic inflammatory disorder of the gastrointestinal (GI) tract. It is believed to be caused by a dysregulation of immune response to pathogens. IBD includes two major forms: ulcerative colitis (UC) and Crohn’s Disease (CD) ^1^. The global prevalence of IBD increased from 3.7 million in 1990 to more than 6.8 million in 2017 ^2^. The detailed etiology of IBD is still unclear, but genetic predisposition, intestinal dysbiosis, various pro-inflammatory cytokines, T-helper cells and interleukins are implicated in the pathogenesis of the disease ^3, 4^. In early stage IBD, a loss of intestinal epithelial barrier function is observed, increasing bacterial translocation which activates the mucosal immune system and intestinal inflammation ^5^. If left unmanaged, patients with UC will have 2% higher incidence of CRC after 10 years, 8% after 20 years and about 18% after 30 years post diagnosis compared to those without IBD and so require frequent surveillance. ^6^. IBD-associated CRC (IBD-CRC) carries a particularly poor prognosis (mortality >50%) ^7^ compared to sporadic CRC, for which prognosis is improving ^8^. Unlike sporadic CRC, which progresses through an adenoma-carcinoma pathway, IBD-CRC evolves from an inflammation-dysplasia-carcinoma sequence ^8^. There are currently no tools that are specifically designed to help clinicians track the initiation and progression of IBD-CRC pathology. Detecting and tracking chronic inflammation in the gastrointestinal (GI) tract is critical to improving outcomes among patients with IBD. Current diagnostic and surveillance methods for IBD are comprised of clinical manifestations (e.g. bloody diarrhea) in conjunction with a physical examination, endoscopy and pathological findings ^9^. Molecular assays measuring elevated biomarkers of inflammation in blood and stool reinforce its evaluation and diagnosis; albeit, no loco-regional information on inflamed areas is acquired ^10^. Currently, endoscopy with biopsy is the gold standard procedure for the diagnosis of IBD and staging the disease activity ^11^. However, endoscopy is invasive and can lead to a toxic megacolon if done during exacerbation. Another drawback of endoscopy is that it is limited to imaging segmental areas of the intestine, cannot provide detailed molecular information regarding the disease and is limited to imaging the superficial mucosal surface ^12^. None of the available standard-of-care diagnostic tools, whether used alone or in combination, completely meet the need for safe, accessible, reliable, quantitative visualization of GI inflammation with high spatial and molecular specificity. Therefore, a novel less invasive and quantitative tool which can provide both functional and morphological information on specific pathologic processes in real time is warranted ^13^.

Numerous studies have explored the utility of [^18^F]-Fluorodeoxyglucose ([^18^F]-FDG) PET in the assessment of IBD. High uptake of [^18^F]-FDG in intestinal tissues due to increased immune metabolic activity allowed delineation of inflammatory sites in IBD patients ^14, 15^. While [^18^F]-FDG has been shown to be sensitive in the detection of IBD, its specificity leaves much to be desired due to physiological uptake in normal bowel ^16^. In recent years, several immunopositron emission tomography (immunoPET) tracers were developed and explored for molecular imaging of chemically induced colitis mouse models using dextran sodium sulfate (DSS) ^17, 18^. Specific immune signatures were targeted for immunoPET imaging. CD4^+^ T cells were detected in colitis mice via a ^89^Zr-labeled diabody specific for the molecule ^17^. Dmochowska and colleagues demonstrated the detection of interleukin-1β, a pro-inflammatory cytokine and CD11b, a marker of neutrophils, dendritic cells and macrophages in DSS-treated mice ^12^. Though a clear advancement in non-invasive imaging for IBD, there are many other pro-inflammatory immune cell phenotypes that contribute to colitis, including neutrophils, dendritic cells, and macrophages, not all of which will be detected using CD4^-^ or CD11b^-^ specific tracers. ImmunoPET using [^89^Zr]Zr-α-IL-1β provides a more tissue-specific signal than the CD11b tracer. However, the role of IL1β as a key driver of inflammation is questionable, as inhibition of IL1β in IBD patients does not generate a positive response ^19^.

The cytokine IL23 was recently recognized for its crucial role in the pathogenesis of IBD ^20, 21^. IL23 is a pro-inflammatory cytokine belonging to IL12 cytokine family. It consists of two subunits, p19 and a p40 subunit shared with IL12 cytokine ^22^. IL23, along with TNFα, triggers a pro-inflammatory cascade that is relevant to IBD pathophysiology. IL23 induces inflammation by promoting the proliferation and differentiation of näive CD4^+^ T cells into pathogenic Th17 cells, which then release other cytokines ^4, 23, 24^. Because of the central role of IL23 in inflammation, it has become a relevant clinical target in IBD treatment. For example, ustekinumab (Stelara™), an IL12/23p40 antagonist is approved by FDA for IBD treatment and clinical data show patients experience symptom relief and achieve clinical remission ^25, 26^. Other drugs targeting IL23 via its p19 subunit are currently in clinical trials^27^. In addition, IL23 is also one of the identified mediators of anti-TNFα drug resistance ^28^, which can only be identified via colonoscopy-obtained tissue biopsies. Taken together, IL23p40 is an appealing target not only for treatment, but also for diagnosis and staging of IBD.

Herein, we present our study wherein we developed an immunoPET imaging agent by radiolabeling a mouse anti-IL23p40 with the radioisotope ^89^Zr (t_1/2_ ∼ 3.27 d). We evaluated the potential of the probe to detect the cytokine in an acute model of gastrointestinal inflammation. We performed competitive binding studies to demonstrate the tracer’s specificity for the cytokine and evaluated whole body tissue distribution to determine its pharmacokinetic parameters. Finally, we correlated the uptake of the radiotracer with serum levels of IL23p40.

## Materials & Methods

### General

All chemicals and supplies were purchased from commercial suppliers and used without further manipulation except when otherwise stated. [^89^Zr]Zr-oxalate was provided via 3D Imaging, LLC (Little Rock, AR). Anti-IL23p40 monoclonal antibody was purchased from Bio X Cell (Cat. No. BE0051, Lebanon, NH, USA).

### Chemical induction of colitis via DSS

All animal experiments were conducted in compliance with the Institutional Animal Care and Use Committee (IACUC) at Wayne State University. Male and female BALB/c mice aged 10-14 wks (20-30 g) were purchased from Charles River Laboratories. We classified the mice based on the gender to evaluate the effect of sex on pathogenesis of IBD and tracer uptake. Colitis was induced by replacing normal drinking water with 3% (w/v) DSS (molecular mass, 40 kDa Alfa Aesar) for 7 days. Mice were assessed and weighed daily for signs of colitis. Severity of colitis was examined by disease activity index (DAI) from Freise *et al* ^*17*^.: weight loss (0, none; 1, 1%–4%; 2, 5%–10%; 3,11%–20%; 4, .20%), fecal blood (0, none; 2, blood present in stool; 4, gross bleeding from anus), and stool consistency (0, normal; 1, moist/sticky; 2, soft; 4, diarrhea).

### Antibody conjugation and ^89^Zr radiolabeling

p-Benzyl-isothiocyanate-deferoxamine (DFO-Bz-SCN, Macrocylics) was conjugated to anti-IL23p40 similar to prior protocols ^29^. Briefly, DFO-Bz-SCN in DMSO was added to the antibody at a ratio of 1:5 (Ab:DFO) in 0.9% saline, pH∼9 at 37 °C for 1 h. Subsequent purification using a centrifugation column filter (MWCO: 30 kDa) and 0.9% saline as the mobile phase eliminated unconjugated DFO-Bz-SCN. ^89^Zr-radioabeling of DFO-anti-IL23p40 antibody proceeded in a neutral pH environment in saline. Approximately 37 MBq (1 mCi) of [^89^Zr]Zr-oxalate previously neutralized to pH 7.0–7.2 was added to a solution of DFO-anti-IL23p40 (0.2 mg, 1.3 nmol) and incubated for 1 h at room temperature. The radiolabeling reaction and efficiency were monitored via radio-instant thin layer chromatography using iTLC-SG-Glass microfiber chromatography paper impregnated with silica gel (Agilent Technologies, Cat. No: SGI0001) as stationary phase (radio-iTLC, Mini-Scan/FC, Eckert and Ziegler) and 50 mM ethylenediaminetetraacetic acid (EDTA) as mobile phase. The reaction was quenched with 5 μL of 50 mM EDTA and purified through spin column centrifugation (MWCO: 30 kDa) with sterile saline as eluent. Stability studies of [^89^Zr]Zr-DFO-anti-IL23p40 in saline at 37 °C were conducted and monitored over 96 h.

### *In vitro* binding and blocking studies

Saturation binding studies were conducted to evaluate the dissociation constant (K_d_) of the [^89^Zr]Zr-DFO-anti-IL23p40. Mouse IL12p40 protein (Bio-Techne MN, USA) (5 µg/ml in bicarbonate buffer) was coated onto 96 well strip plate and incubated at 4 °C overnight. The wells were then washed 3× with 0.05% Tween-20 in 1× PBS and incubated for 2 h at room temperature with 200 µL of blocking buffer (1% BSA in wash buffer). The wells were washed and incubated with varying concentrations of radiolabeled antibody (0.5 – 5000 nM) in triplicate. After 1 h at 37 °C, the unbound radiotracer was removed and the bound activity for each well was measured by a gamma counter. The dissociation constant, K_d_, was calculated by nonlinear regression using GraphPad Prism v. 9.2.

To evaluate specific binding, a separate group of wells were treated with excess concentration of cold mAbs (500 nM) 30 min prior to the addition of the radiotracer (5 nM, 0.34 μCi, 0.012 MBq per well). Specific binding was measured by subtracting nonspecific binding from total bound activity.

### PET imaging

[^89^Zr]Zr-DFO-anti-IL23p40 (6.66-9.25 MBq, 36-50 ug, 0.24-0.33 nmol) in sterile saline was administered intravenously (i.v.) in mice (n=5 per sex) on the lateral tail vein five days after DSS treatment. Imaging experiments were conducted using Bruker Albira Si PET from 24 to 96 h post-injection (p.i.) while the mice were anesthetized with 2% isoflurane. Images were reconstructed through Maximum Likelihood Expectation Maximization (MLEM) in 12 iterations and 0.75 mm voxel resolution and were analyzed by PMOD version 4.3 software. Volumes-of-interest (VOI) were acquired via isocontouring and expressed as the mean percentage of injected dose per mL of tissue (%ID/mL). A separate cohort of DSS-treated mice were injected with an ^89^Zr-labeled nonspecific isotype-matched IgG antibody to control for specificity. Images were then acquired 48 h p.i. After imaging, mice were euthanized, and the colons were excised and laid out for *ex vivo* PET imaging.

### Biodistribution and *in vivo* blocking studies

The tissue distribution of [^89^Zr]Zr-DFO-anti-IL23p40 was assessed in healthy and colitis mice by i.v. injection of 0.37-0.92 MBq (10−25 µCi, 2-5 μg) on the lateral tail vein of the mice. Euthanasia was performed 48 h p.i. via CO_2_ asphyxiation. Blood was immediately collected via cardiac puncture. Select organs were collected and weighed. Bound radioactivity was measured using a gamma counter (Perkin Elmer 2480 Wizard2). To assess specificity of target binding, we performed an *in vivo* blocking study where 100× of unlabeled antibody was co-injected with the radiotracer in a separate group of mice.

### IL23p40 ELISA

Blood was collected via the cheek vein prior (baseline or day 0), during (day 5) and after (day 8) DSS treatment. Sera were obtained and stored at -80 ºC to allow for decay prior to analysis. Levels of IL23p40 were evaluated using a commercial ELISA kit (RayBiotech, Cat. No. ELM-IL12p40p70).

### In Situ Hybridization

To investigate the expression of IL23p40 in colitis, RNA *in situ* hybridization for IL23p40 mRNA (*IL23b*) was performed using an RNAscope 2.5 HD (ACD Bio) detection assay. RNA spatial expression of IL12 (*IL12a*) and IFN-γ (*IFNg*) were also examined. In brief, intestinal tissues were harvested, fixed, and stored in 10 % formalin until decayed. Once decayed, formalin was removed and replaced with 70% ethanol prior to paraffin embedding and sectioning. Tissues were sliced into 5 µm sections and mounted on glass slides. Paraffin-embedded tissue sections were deparaffinized and were subjected to antigen retrieval using RNAscope® antigen retrieval kit (ACD Bio) followed by hybridization, signal amplification and chromogenic detection procedures which were performed according to the manufacturer’s instructions using RNAscope® probe Mm-IL12b (ACD Bio), Mm IL12a (ACD Bio) and RNAscope® 2.5 HD duplex detection reagents (ACD Bio). IL23 and IL12 were detected using red and green chromogen respectively. The images (40× objective) were captured under a light microscope (Carl Zeiss) and signal quantification was analyzed by HALO software (Indica Labs).

### Statistical analysis

GraphPad Prism version 9.02 was used to perform statistical analyses unless otherwise stated. Data are presented as the mean ± S.D. Mann-Whitney t-tests were carried out for tumor blocking competitive studies. Two-way ANOVA multiple comparison analyses were performed on tissue distribution studies. The significance of correlations was determined by Pearson analysis. A P value < 0.05 was considered statistically significant.

### Study approval

All animal experiments were conducted in compliance with the Institutional Animal Care and Use Committee (IACUC) at Wayne State University.

## Results

### Successful labeling and target binding of [^89^Zr]Zr-DFO-anti-IL23p40

Radiolabeling of the DFO immunoconjugate was straightforward with radiolabeling yields of >95 % obtained. A specific activity of 180 ± 10 MBq/mg was established. The radiotracer remains moderately intact (>90%) in saline over a 96-h incubation period (Supplementary Fig. S1). The binding of [^89^Zr]Zr-DFO-anti-IL23p40 to mouse IL23p40 protein was evaluated by saturation binding assays, which confirmed preserved high affinity (K_d_ of ∼ 9.8 ± 1.9 nM) of the antibody for its target (Fig. 1A). The *in vitro* blocking study incubating [^89^Zr]Zr-DFO-anti-IL23p40 with 100-fold mass excess of the unmodified non-radiolabeled antibody demonstrated significantly decreased binding of [^89^Zr]Zr-DFO-anti-IL23p40 to IL23p40 (P < 0.0001) (Fig. 1B). These findings confirm the specificity of the radiolabeled antibody to IL23p40 protein *in vitro*.

**Figure 1.**
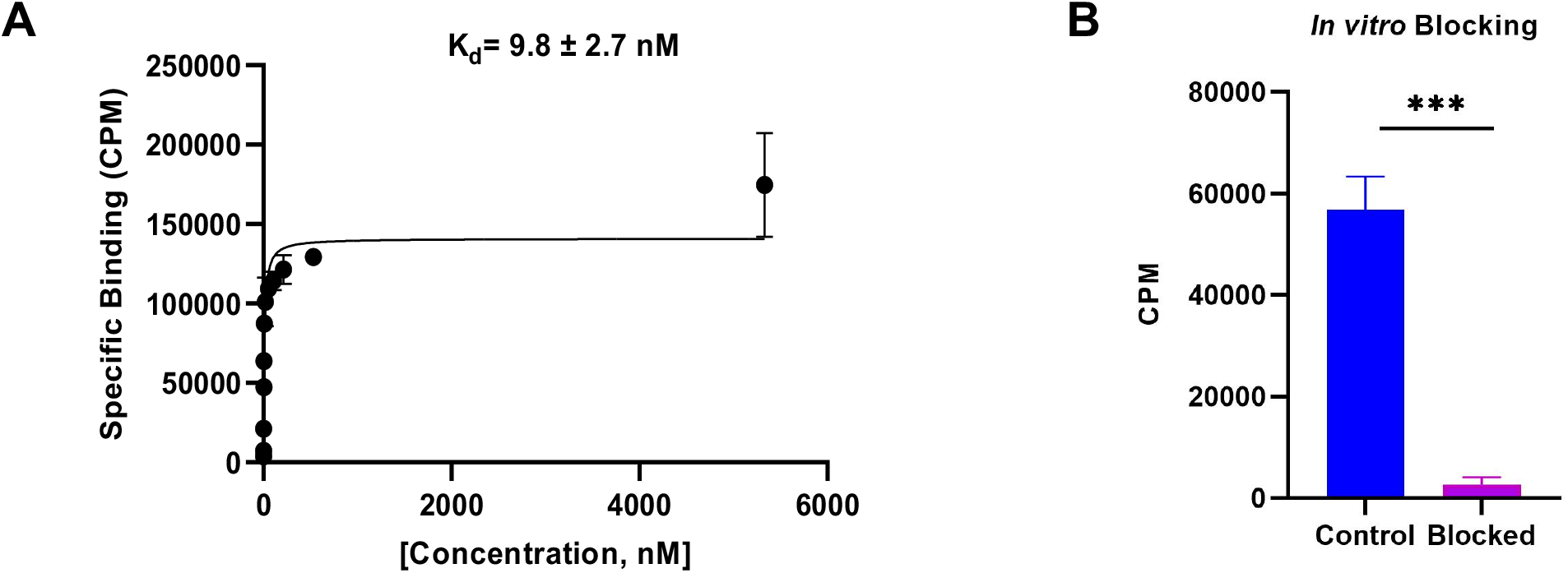
*In vitro* characterization of [^89^Zr]Zr-DFO-anti-IL23p40. (A) Non-linear regression analysis determined the binding affinity expressed as K_d_ of the tracer for IL23p40. (B) Competitive inhibition with 100× of unmodified antibody co-administered with the radiotracer in IL23p40-coated wells.

### DSS-induced colitis

Colitis was induced in Balb/c mice by giving DSS *ad libitum* for 7 days followed by recovery with regular drinking water on day 8 (Fig. 2A). DSS treated mice showed weight loss beginning on day 5 (Fig. 2B). DAI scores for stool consistency and fecal bleeding were higher in the DSS-treated mice compared to control mice. DAI score in the most DSS treated mice at day 7 was 3 for stool consistency (diarrhea) and 2 for fecal blood (blood present in stool).

**Figure 2:**
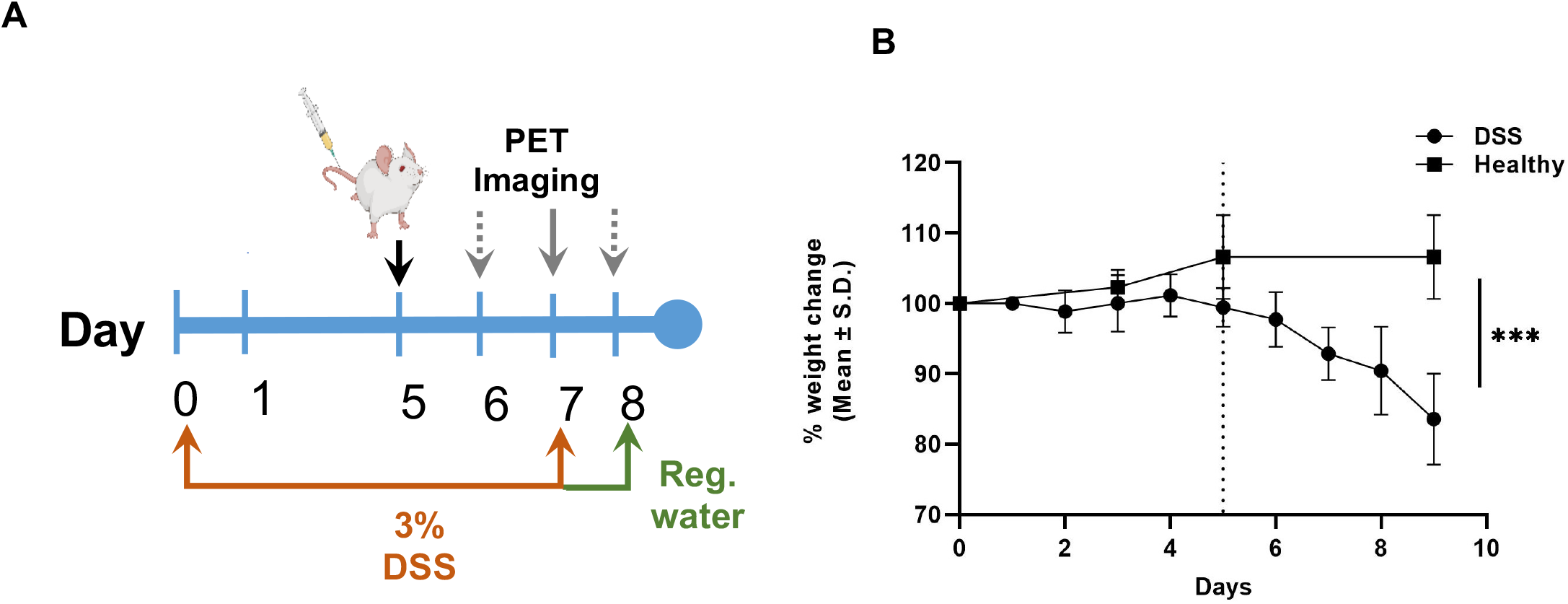
(A) Scheme showing days of DSS treatment, injection of [^89^Zr]Zr-DFO-anti-IL23p40 and PET imaging. (B) Change in body weight was observed after five days of DSS treatment. The vertical dashed line indicates the beginning of body weight lost.

### *In vivo* delineation of intestinal inflammation by IL23p40 immunoPET

[^89^Zr]Zr-DFO-anti-IL23p40 delineated the large intestinal tract of DSS-induced colitis mice whereas no clear delineation was observed in healthy control mice (Fig. 3A). Time activity curve of [^89^Zr]Zr-DFO-anti-IL23p40 examines heart, liver, and muscle uptake from 24 to 96 h showed decreasing non-specific binding of radiotracer over time. (Fig. 3B). Uptake of the radiotracer in the colons of both healthy and DSS-treated mice were obtained and classified according to sex (Fig. 3C-D). For both male (DSS: 5.99 ± 0.65 %ID/mL vs. Control: 3.98 ± 0.58 %ID/mL, p<0.0018) and female mice (DSS: 8.53 ± 1.47 %ID/mL vs. Control, 6.56 ± 0.85 %ID/mL, p<0.02), DSS treatment led to higher uptake in the colon than the control group at 24 h p.i. The specificity of the cytokine radiotracer was assessed by imaging a separate cohort of DSS-treated mice with a ^89^Zr-radiolabeled isotype control antibody. Unlike colitis mice imaged with [^89^Zr]Zr-DFO-anti-IL23p40, colitis mice imaged with [^89^Zr]Zr-DFO-IgG did not show specific accumulation in the intestines (Fig. 3E-G). We next performed *ex vivo* imaging to assess for specific focal uptake of the tracer in colons. *Ex vivo* PET imaging showed increased uptake in the cecum, proximal and mid colon of colitis mice injected with [^89^Zr]Zr-DFO-anti-IL23p40 whereas lower accumulation of [^89^Zr]Zr-DFO-IgG was observed in similar regions (**Fig. 4A**).

**Figure 3:**
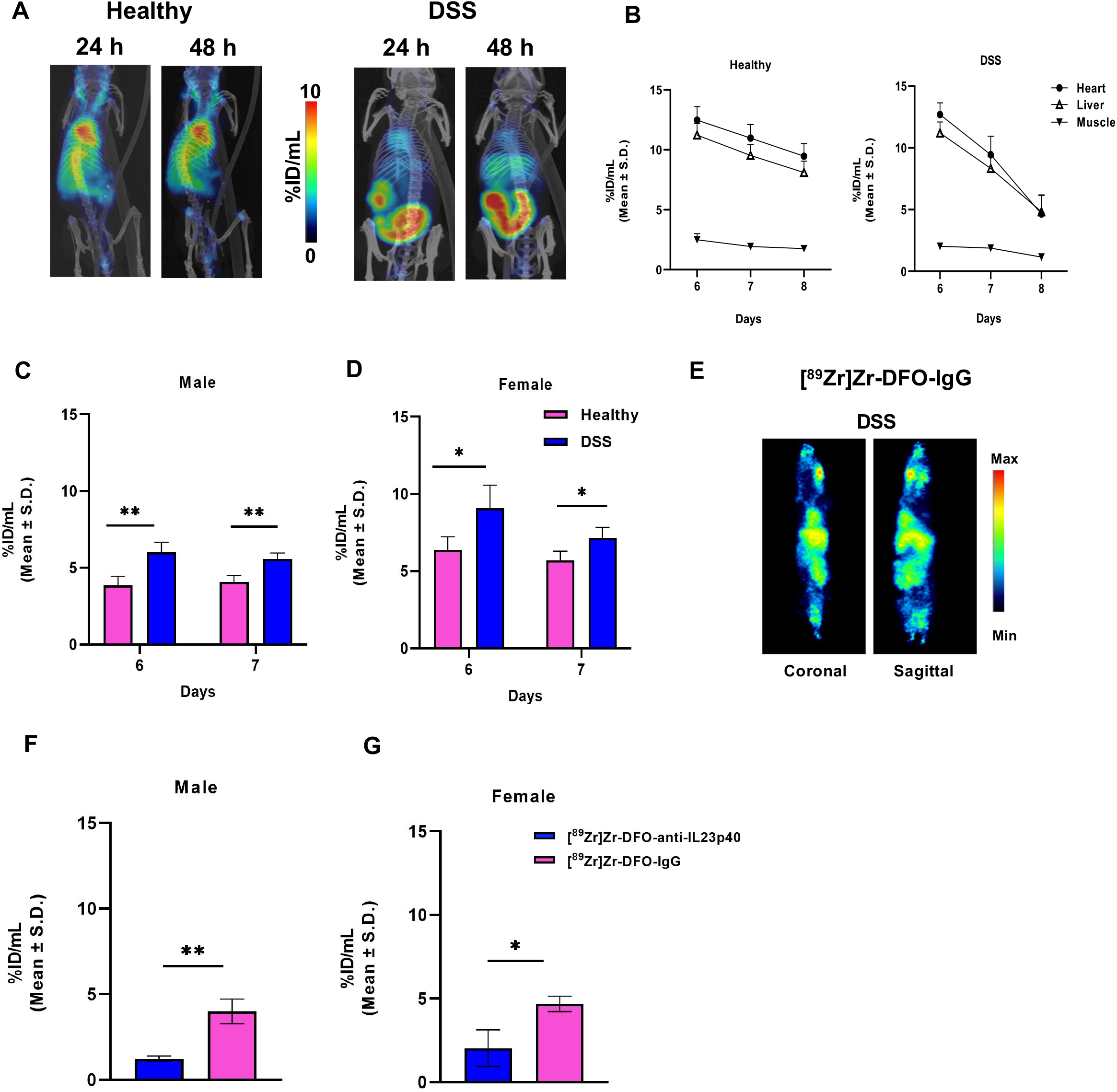
Immuno-PET imaging of healthy and DSS-treated mice. (A) Representative PET/CT (coronal and sagittal) images of control (left) and DSS-treated mice (right) acquired 24 and 48 h after injection of the radiotracer. (B) Time activity curves display of radiotracer binding in heart, liver and muscle in healthy and DSS treated group. (C) Accumulation of the radiotracer in colon in male (D) and female. (E) Representative PET image of a DSS-treated mouse after injection with [^89^Zr]Zr-DFO-IgG. (F). VOI of colon in DSS treated mice for [^89^Zr]Zr-DFO-anti-IL23p40 and [^89^Zr]Zr-DFO-IgG in male and (G) female group. *P < 0.05. **P < 0.01. ***P< 0.001. **** P<0.0001.

**Figure 4:**
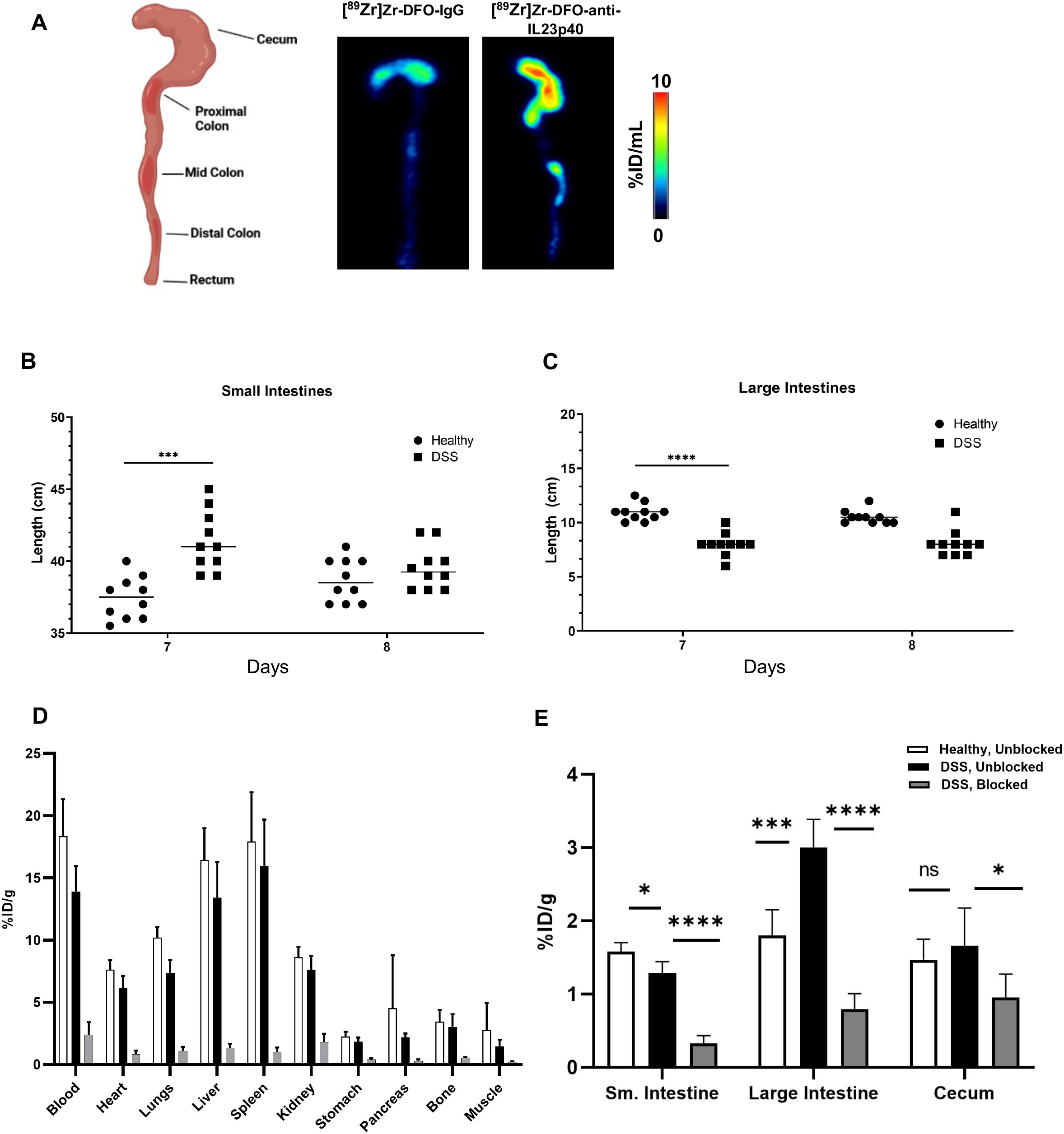
*Ex vivo* immuno-PET and tissue distribution studies. (A) Representative PET images of excised intestinal tissues of DSS-administered mice demonstrate higher accumulation of the IL23p40 tracer (middle) in the cecum, proximal and mid colon versus a radiolabeled non-specific isotype antibody (right). Scheme of a mouse colon depicting colon sections (left) (Created via Biorender). Lengths of (B) small and (C) large intestines from separate cohorts were measured on the last day of DSS treatment (day 7) and two days after switching to regular drinking water (day 9). (D) Tissue distribution of the IL23p40 radiotracer in control and DSS treated mice at 48 h p.i. A separate DSS-treated cohort was co-injected with excess unmodified anti-IL23p40 for competitive binding with the radiotracer. (E) Uptake of the tracer in small intestines, colon and cecum. *P < 0.05. **P < 0.01.

The length of the small (**Fig. 4B**) and large intestines (**Fig. 4C**) were measured post-imaging. The small intestines of DSS treated mice appeared longer than healthy mice while the large intestines are collectively shorter. No difference was displayed in intestinal lengths of the mice who were switched to regular drinking water after acute DSS administration, suggesting quick recovery in these groups.

### Tissue biodistribution study

Tissue biodistribution of [^89^Zr]Zr-DFO-anti-IL23p40 was performed in control and DSS-treated mice at 48 h p.i (**Fig. 4D**). Uptake in the colon was higher for the DSS group than for the controls (3.0 ± 0.4 %ID/g vs. 1.7 ± 0.35 %ID/g, P<0.01). No difference was observed in the small intestines. The radiotracer uptake in other healthy tissues was comparable.

To confirm specificity, blocking experiments were performed wherein at least 100-fold excess of the unmodified, non-radioactive antibody was co-injected with the radiotracer in DSS-treated mice. A notable decrease in accumulation of the radiotracer was observed in tissues of the blocked cohort was observed versus unblocked colitis cohorts and healthy controls (**Fig. 4D**). Blocking significantly decreased the uptake in small intestines (1.3 ± 0.2 %ID/g vs. 0.4 ± 0.1 %ID/g, P<0.0001), colon (3.0 ± 0.4 %ID/g vs. 0.8 ± 0.2 %ID/g, P<0.0001) and cecum (1.6 ± 0.5 %ID/g vs 1.0 ± 0.3 %ID/g, p <0.01) (**Fig. 4E**).

### Correlation between tracer uptake and serum expression of IL23p40

To confirm expression of IL23p40 in colitis mice, sera were sequentially collected at baseline prior to DSS treatment (Day 0), on day 5 before radiotracer injection. ELISA results showed elevated total IL23p40 on day 5 from baseline (**Fig. 5A**) (Day 5: 2.25 ± 0.63 vs. baseline: 1.53 ± 0.46 ng/mL p=0.03). A correlation between IL23p40 levels in serum on day 5 and [^89^Zr]Zr-DFO-anti-IL23p40 uptake in the small intestine and cecum indicate a direct relationship between IL23p40 concentration and radiotracer uptake (**Fig. 5B**). A positive trend which was observed with cytokine expression in the serum and tracer uptake in colon was not significant (**Fig. S2**).

**Figure 5:**
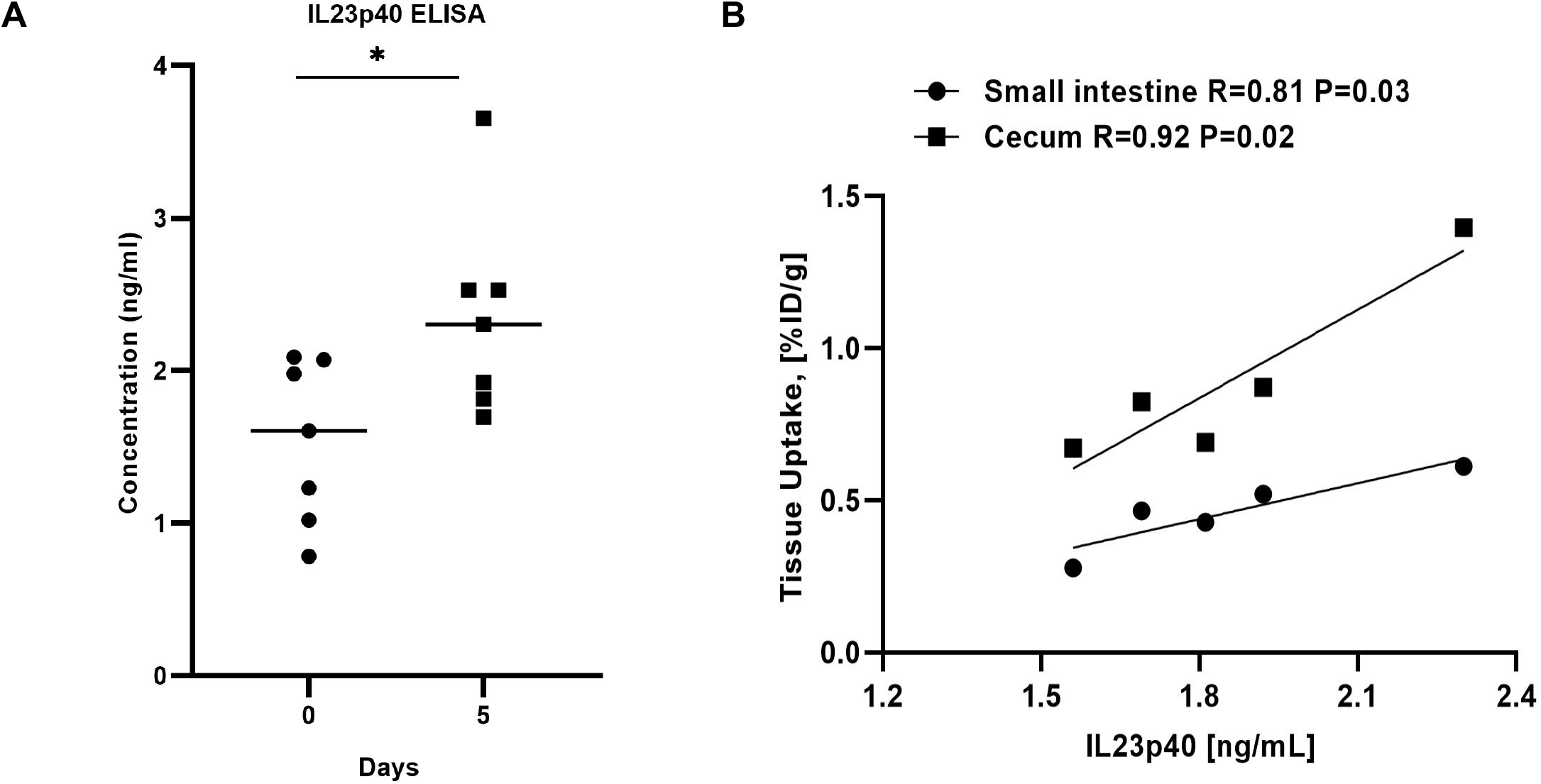
Correlation of serum IL23p40 with [^89^Zr]Zr-DFO-anti-IL23p40 tissue uptake. (A) IL23p40 in serum showed increased expression five days after DSS administration. (B) A strong positive correlation between IL23p40 serum concentration and radiotracer uptake is demonstrated.

### High transcript expression of IL23 in intestinal tissues confirms the PET images

To further detect IL23p40 cytokine in healthy and DSS mice, *in situ* hybridization was performed to visualize single RNA molecules per cell. IL12b, the gene for IL23p40, was highly expressed in the cecum of DSS-administered mice versus control (**Fig. 6A**). A high percentage of positive cells (red positive cells / total cells) demonstrated significant expression of *IL12b* in the DSS group (**Fig. 6B**) compared to control. Transcripts of IL12p35 were not as markedly expressed as IL23 (IL12b) in inflamed tissues (**Fig. S3**), suggesting that the driver of inflammation in this setting is IL23. Interestingly, IFN-γ appears to be expressed in the DSS-treated intestines (**Fig. S3**).

**Figure 6:**
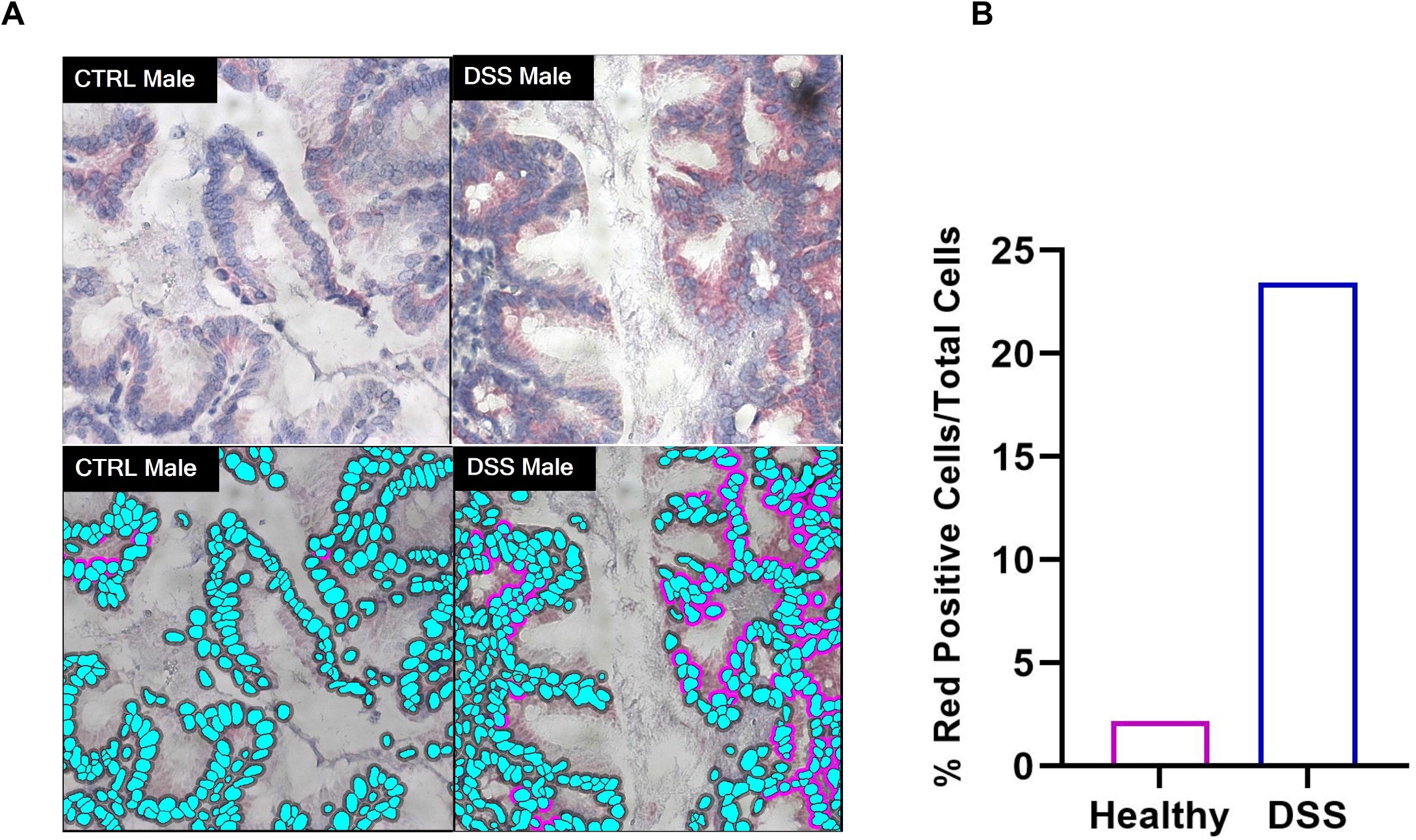
*In situ* hybridization to detect expression of IL23 in intestinal tissues of control and DSS mice. (A) IL23 was detected using red chromogen. Higher red signal which indicated IL12b gene is observed in DSS treated mice (top left) compared to healthy mice (top right). Bottom figures represent individual cells provided by HALO software. The green color indicates hematoxylin staining of cell nuclei. The purple color are the positive cells containing red signal. (B) The quantitative analysis of the mRNA levels of IL23 based on the percentage of positive cells for red signal.

## Discussion

IBD is anticipated to impose increasing health and economic burdens as it appears to be rising globally ^2^. Patients from industrialized western countries are primarily afflicted with IBD. Albeit, increasing prevalence and incidence are observed worldwide, specifically in newly industrialized countries ^2^. In the US alone, approximately 3.1 million adults were diagnosed with this autoimmune condition in 2015 with increased prevalence in >45 years old, Hispanic and non-Hispanic white people and those with low level of education ^30^. Thus, addressing this autoimmune condition is critical before it becomes a global issue. In addition, management and surveillance of IBD is crucial as it predisposes patients to a higher risk of developing colorectal cancer (CRC) ^31^. Colitis is further observed as an immune-related adverse event associated with Immune checkpoint inhibitors (ICIs) ^32^, which can be life-threatening at higher grades ^33^. Current diagnostic standard-of-care is reliant on endoscopy or colonoscopy procedures to assess mucosal inflammation. However, as mentioned above, endo/colonoscopy can only image superficial tissue architectures. These approaches reveal very little about the molecular characteristics of a patient’s disease especially when inflammation is occurring deep within the mucosal layer before anatomical changes are observed ^34^. Moreover, this approach carries a risk of bowel perforation that can lead to bleeding. Non-invasive molecular imaging techniques can potentially overcome these limitations in current standard of care and improve IBD diagnosis.

Herein, we laid down the development of a non-invasive and quantitative imaging agent via immunoPET that is directed against IL23p40 for detection of acute inflammation in DSS mouse model of colitis. In this study, we have shown that [^89^Zr]Zr-DFO-anti-IL23p40 was able to detect IL23p40 in the colons of colitis mice. We have demonstrated the radiotracer’s specificity in both *in vitro* and *in vivo* competitive binding studies as well as when compared against a radiolabeled non-specific antibody, which failed to delineate intestinal inflammation. Results of the biodistribution study clearly displayed increased uptake in gastrointestinal tissues of the colitis mice, reflecting the PET imaging results. In our hands, intestinal inflammation developed five days post-administration of DSS as marked by the significant weight loss in mice and increased disease activity index score. Different small and large intestinal lengths in healthy versus DSS mice further confirm the gastrointestinal inflammatory status of the latter with measurements in good agreement with prior reports ^17^. [^89^Zr]Zr-DFO-anti-IL23p40 was injected on this day (day 5) with the optimum imaging time point occurring between 24 and 48 h p.i. where contrast is high.

Growing lines of evidence have demonstrated that circulating IL23 may be a biomarker of disease severity in patients with IBD^35^. Lucaciu et al. have shown that IL23 (from serum) is a superior diagnostic marker for differentiating mild versus severe inflammation than current standard-of-care inflammation markers like C-reactive protein and fecal calprotectin ^36^. In our study, elevated IL23 was detected by the radiotracer as evidenced by a positive correlation with serum IL23p40. Thus, the utility of IL23 PET imaging to quantitatively evaluate the severity of the inflammatory disease via IL23 PET imaging will be done in future studies to establish its diagnostic potential.

The gender disparity in inflammatory response to mucosal injury was evident in our study and was clearly delineated by the radiotracer. DSS induced colitis in male mice had significantly higher inflammation and crypt damage than in female rodents, which parallels findings by others ^37^. Sex differences have long been recognized in IBD susceptibility and disease severity, but are not fully understood ^38^. The estrus cycle has been shown to influence IBD pathogenesis by controlling intestinal permeability; permeability is lower at the pro-estrus stage, when estrogen is at peak^39^. Homma *et al*. has shown that the gut of female rodents withstood the damaging effects of hypoxia and acidosis versus those of male intestines but was reversed upon administration of estradiol to male rats; thus, demonstrating a protective role of estrogen^35^. Moreover, estrogen was proven to play an anti-inflammatory function in the gut by reducing immune infiltration and cytokine secretion during colitis^40, 41^, which led to significantly lower inflammation as demonstrated by Harnish et al.

Other molecular imaging agents are investigated to diagnose IBD. To date, clinical SPECT and PET imaging of IBD is restricted to using ^111^In or ^99m^Tc-radiolabeled leukocytes and [^18^F]-FDG, none of which provides information on specific immune mediators ^15, 42^. Leukocyte SPECT imaging suffers from poor quality images with the radiolabeling a time-consuming process ^43^. [^18^F]-FDG PET can identify inflammation but its highly variable tissue uptake limits its utility as a diagnostic tool ^44^. FDG marks cellular metabolism, providing information about the energy consumption of mucosal layer infiltrating immune cells within the inflamed tissue. This can be useful for imaging glycolytic activity of immune cells, but not for assessing mediators of inflammation ^14^. New immunoPET agents were developed and has shown excellent target selectivity; however, to-date, its use for IBD detection is limited to preclinical studies. Freise *et al*. showed delineation of CD4^+^ T cells that are present in inflamed colons of DSS mice using a murine CD4 diabody (GK1.5) radiolabeled with zirconium-89 ^17^. Innate immune markers CD11b and IL-1β were detected by ^89^Zr-conjugated antibodies targeting these molecules. Both immunoPET agents detected colonic inflammation but [^89^Zr]Zr-α-IL-1β was more specific in accumulation in gastrointestinal tract compared to [^89^Zr]Zr-α-CD11b which was dispersed to other tissues ^12^. Both immunoPET agents were able to reliably detect inflamed colons. However, imaging CD4 precludes assessment of other pro-inflammatory immune cell phenotypes that contribute to this autoimmune condition. On the other hand, the role of IL-1β in IBD is far from clear, stemming from the lack of positive response in patients given the targeted blockade ^12^.

Our previous work developed and assessed an immunoPET imaging agent specific for IL12p70, which detects the cytokine globally in an induced inflammation model ^45^. However, compelling studies have shown that IL23 – not IL12 plays a major pro-inflammatory role in IBD ^21^. The FDA approval of ustekinumab (Stelara®), an anti-IL23p40 monoclonal antibody for treatment of CD patients and those with moderate to severe UC further supported our rationale toward the development of a companion diagnostic that targeted the cytokine. The main limitation of this study lies in the fact that the tracer maybe detecting IL12, as it shares the p40 subunit with IL23. Nevertheless, IL23 (*IL12b*) is abundantly produced than IL12 (*IL12a*) in our mouse model as demonstrated by RNA *in situ* hybridization (Fig. 6 and Fig. S3). Another limitation lies in the potential of the antibody to neutralize IL23p40, consequently decreasing IL23 expression which we have observed during the *in vivo* competitive binding studies as well as during imaging studies. To address this, work is underway to develop a non-neutralizing antibody carrier specific to IL23.

Our finding validates our IL23p40 tracer as a novel approach to delineate IBD selectively and robustly in this model of acute inflammation. The promising findings from this study have major implications in IBD standard-of-care as it demonstrated proof-of-concept for a non-invasive, quantitative and molecular technique for diagnosis of IBD. Furthermore, IL23p40 immunoPET can provide useful insight for patient treatment, including those who potentially respond to IL23 blockade.

## Conclusion

To the best of our knowledge, this is the first report illustrating the successful development of an IBD immunoPET imaging tool specific for IL23p40. This new imaging technology can potentially facilitate early detection and accurate staging of IBD in patients via generation of a global, *in situ* inflammation “map” of the entire gastrointestinal tract.

## Abbreviations used in this paper

IBD: inflammatory bowel disease
CD: Crohn’s Disease
UC: ulcerative colitis
CRC: colorectal cancer
PET: positron emission tomography
IL: interleukin
GI: gastrointestinal
[^18^F]-FDG: [^18^F]- Fluorodeoxyglucose
DSS: dextran sodium sulfate
TNFα: tumor necrosis factor alpha
iTLC: instant thin layer chromatography
EDTA: ethylenediaminetetraacetic acid
VOI: volumes-of-interest

## Acknowledgments

We thank Dr. Lisa Polin and Michael Bradley for suggestions and advice on animal studies. We are further grateful to Dr. Freddy Escorcia for helpful discussions and review of the manuscript. We also thank Dr. Todd Sasser for assistance in the PET/CT image analysis.

## Notes

**Grant support:** The study was supported by NIH/NCI R37 CA220482 (NTV). The Microscopy, Imaging and Cytometry Resources Core (MICR) and Animal Modeling and Therapeutics Core, which provided technical assistance are supported, in part, by the NIH Cancer Center grant P30 CA022453 to the Karmanos Cancer Institute at Wayne State University, the Perinatology Research Branch of the National Institutes of Child Health and Development at Wayne State University.

### Competing Interest Statement

The authors have declared no competing interest.

