## Supplementary Data for "Detection of IL23p40 via Positron Emission Tomography Visualized Inflammatory Bowel Disease"

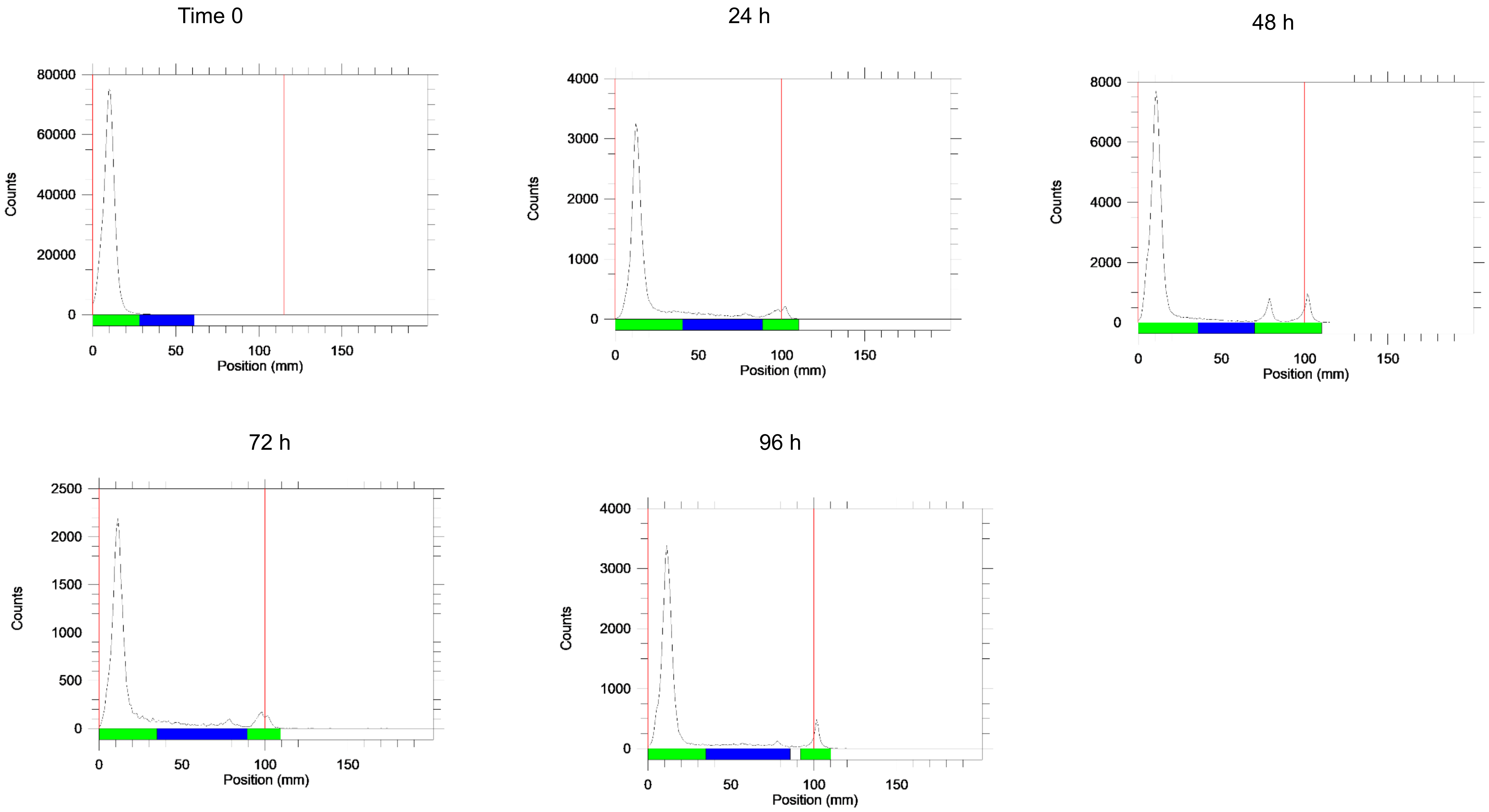


**Figure S.1. Stability of [$^{89}$Zr]Zr-DFO-anti-IL-12/23p40 over time measured by radio-iTLC.**

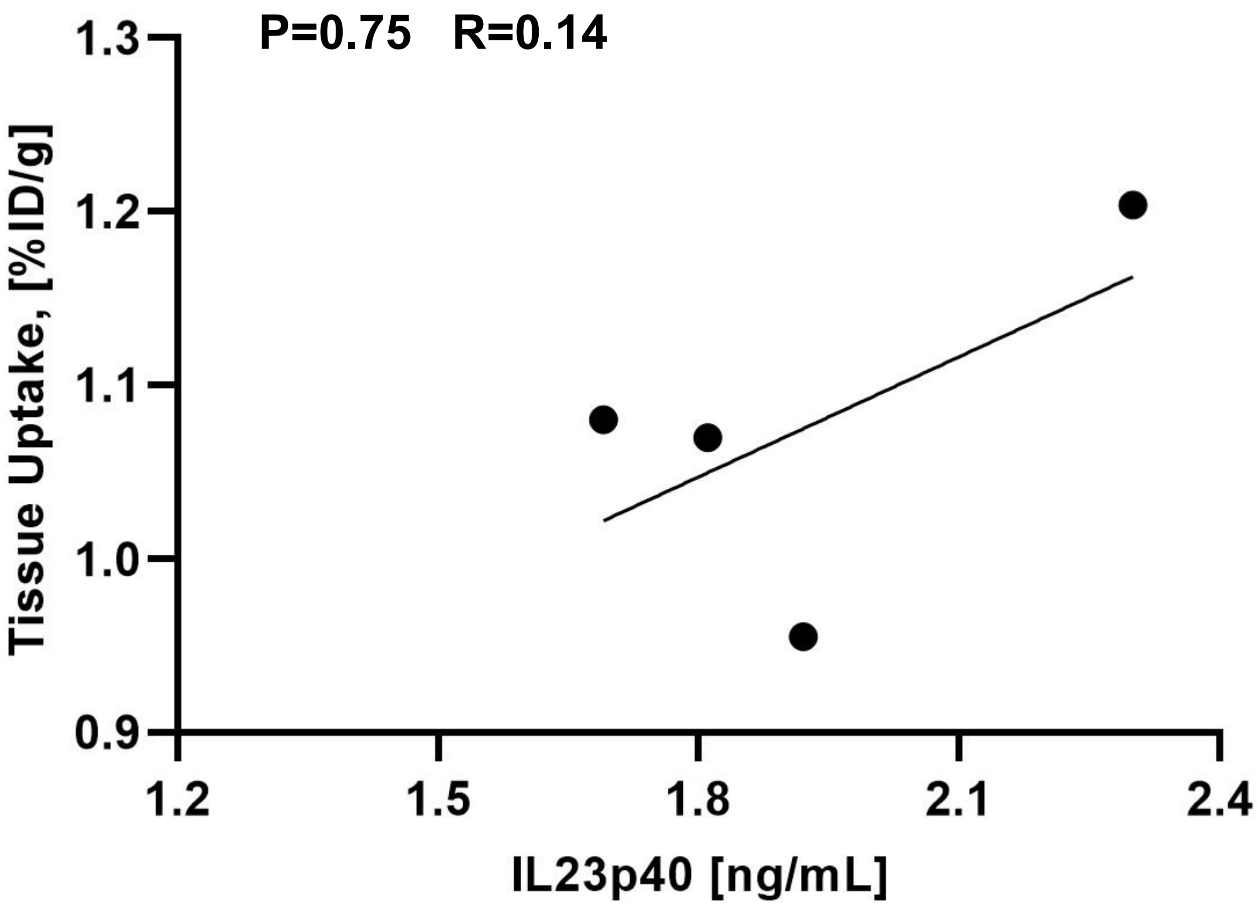


**Figure S.2. Correlation between IL23p40 Concentration in serum and uptake of radiotracer in colon from biodistribution study.**

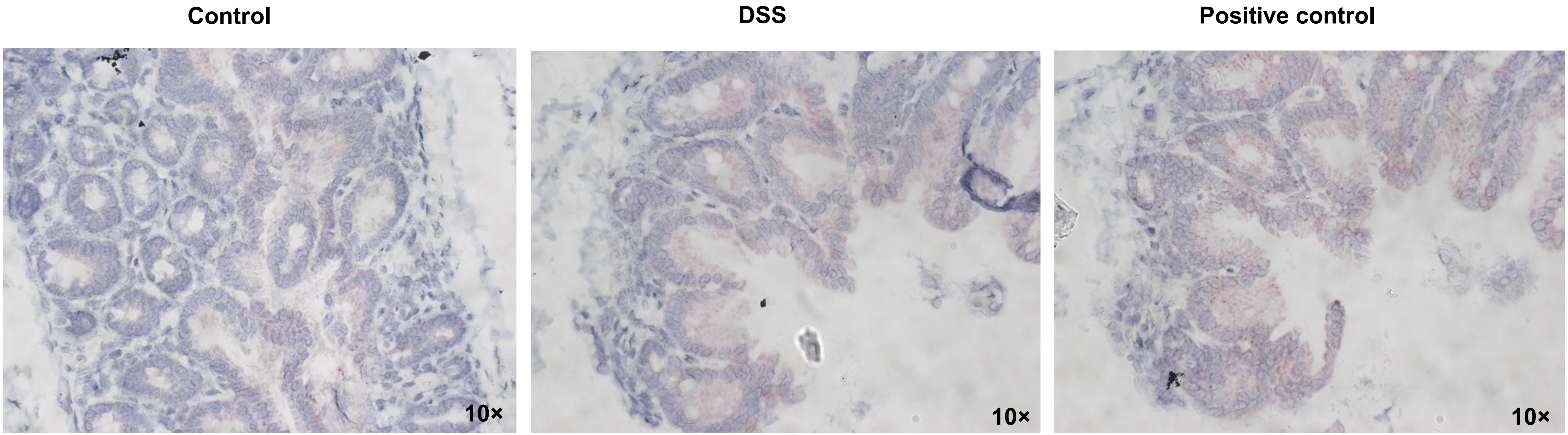


**Figure S.3. RNAscope ISH results for IL12p40 and IFN-γ in control (left), colitis mice (middle) and positive control probe (right). IL12p40 and IFN-γ was detected using green and red chromogen respectively. Purple dots indicate cell nucleus.**
